# Ecological niche modeling the potential geographic distribution of four *Culicoides* species of veterinary significance in Florida

**DOI:** 10.1101/447003

**Authors:** Kristin E. Sloyer, Nathan D. Burkett-Cadena, Anni Yang, Joseph L. Corn, Stacey L. Vigil, Bethany L. McGregor, Samantha M. Wisely, Jason K. Blackburn

## Abstract

Epizootic hemorrhagic disease (EHD) is a viral arthropod-borne disease affecting wild and domestic ruminants. EHD virus (EHDV) is transmitted to vertebrate animal hosts by biting midges in the genus *Culicoides. Culicoides sonorensis* Latreille is the only confirmed vector of EHDV in the United States but is considered rare in Florida and not sufficiently abundant to support EHDV transmission. This study used ecological niche modeling to map the potential geographical distributions and associated ecological variable space of four *Culicoides* species suspected of transmitting EHDV in Florida, including *Culicoides insignis, Culicoides stellifer, Culicoides debilipalpis* and *Culicoides venustus.* Models were developed with the Genetic Algorithm for Rule Set Production in DesktopGARP v1.1.3 using species occurrence data from field sampling along with environmental variables from WorldClim and Trypanosomiasis and Land use in Africa. For three *Culicoides* species (C. *insignis, C. stellifer* and *C. debilipalpis*) 96 – 98% of the presence points were predicted across the Florida landscape (63.77% – 72.53%). For *C. venustus*, models predicted 98.00% of presence points across 27.42% of Florida. Geographic variations were detected between species. *Culicoides insignis* was predicted to be restricted to peninsular Florida, and in contrast, *C. venustus* was predicted to be primarily in north Florida and the panhandle region. *Culicoides stellifer* and *C. debilipalpis* were predicted nearly statewide. Environmental conditions also differed by species, with some species’ ranges predicted by more narrow ranges of variables than others. The Normalized Difference Vegetation Index (NDVI) was a major predictor of *C. venustus* and *C. insignis* presence. For *C. stellifer*, Land Surface Temperature, Middle Infrared were the most limiting predictors of presence. The limiting variables for *C. debilipalpis* were NDVI Bi-Annual Amplitude and NDVI Annual Amplitude at 22.45% and 28.09%, respectively. The model outputs, including maps and environmental variable range predictions generated from these experiments provide an important first pass at predicting species of veterinary importance in Florida. Because EHDV cannot exist in the environment without the vector, model outputs can be used to estimate the potential risk of disease for animal hosts across Florida. Results also provide distribution and habitat information useful for integrated pest management practices.

## Introduction

Vector-borne pathogens can only exist in a permissive environment that supports appropriate vectors (and hosts), such that the distribution of disease is linked to vector distribution. Species distribution models (SDMs) can be used to map the potential distribution of disease vectors allowing inference of disease risk [1–4]. In disease ecology, SDMs are useful to determine the potential current and future geographic distribution of vector species as proxies for the pathogens they transmit [2,4–6]. Modeling probable occurrence is important because the act of *in situ* surveillance requires extensive resources (e.g. personnel, sampling equipment, and laboratory resources) [7]. More simply, map predictions help researchers better understand where vector-borne disease risk is most likely and to target surveillance to areas of highest risk.

Ecological niche models (ENMs) are SDMs commonly used to predict the geographic distribution of a species by determining the most likely environmental conditions associated with collection locations of the target species [8]. The modeling process is rooted in ecological niche theory, with a focus on the Hutchisonian n-dimensional hyper volume [9] of conditions supporting a species on the landscape. A niche is a specific set of environmental conditions which allows for the presence of a species. Simply put, niche theory states that no two species can occupy the same ecological niche, in the case of this work, no two species will cohabitate the same median ranges for all variables. Broadly, ENMs apply each unique set of ecological parameters allowing for a species to maintain a population without immigration [9,10], with a focus on abiotic and climatological conditions that support a species [11]. In general, ENMs use either presence-only or presence and absence data of a target species and environmental covariates to identify non-random patterns relating species occurrence to the landscape [12]. A list of commonly used models include machine learning techniques such as MAXENT [13] Boosted Regression Trees [14], Random Forest [15], and the Genetic Algorithm for Rule-Set Production (GARP) [1,12].

Epizootic hemorrhagic disease virus (EHDV) and blue tongue virus (BTV) (Reoviridae: *Orbivirus*) are economically important pathogens transmitted by biting midges of the genus *Culicoides* Latreille (Diptera: Ceratopogonidae), which can cause disease in a variety of ruminant species worldwide. While many vector species have been identified, vectors have yet to be determined in several regions. In Europe, Asia, and Africa, confirmed BTV vectors and suspected EHDV vectors are *Culicoides obsoletus* (Meigen), *Culicoides scoticus* Downes & Kettle, *Culicoides pulicaris* (Linnaeus), *Culicoides imicola* Keiffer, with *Culicoides dewulfi* Goetghebuer, and *Culicoides chiopterus* (Meigen) also known to transmit the viruses in Europe [16]. In North America, the sole confirmed vector of EHDV is *Culicoides sonorensis* Latreille, which along with *Culicoides insignis* Lutz is also a confirmed vector of BTV [17,18]. This poses an issue for parts of the United States, such as Florida, where EHDV epizootics occur but *C. sonorensis* is absent. Suspected vectors of EHDV in Florida include *C. insignis, Culicoides stellifer* (Coquillett), *Culicoides venustus* Hoffman, and *Culicoides debilipalpis* Lutz [18–21].

Several studies in Europe, Africa, and the Americas have investigated the environmental factors associated with the geographic distributions of multiple *Culicoides* species. In Portugal, generalized linear mixed models were used to demonstrate the most important environmental factors predicting the distribution of European disease vectors [22]. Diurnal temperature range, number of frost days, cold stress, dry stress, and median monthly temperature were the most important factors driving the distribution of *C. imicola* [22,23], the primary BTV vector in Europe. In contrast, the distributions of species within the Obsoletus group (*C. chiopterus, C. dewulfi, C. scoticus, C. obsoletus*, and *Culicoides montanus* Shakirzjanova) were predicted by diurnal temperature range, and median monthly temperature [22]. In North America, Zuliani et al. [24] used a maximum entropy (MAXENT) approach to ENM and temperature variables to predict future distributions of *C. sonorensis* in the western US, due to climate change, and found that northern latitudinal limits are most at risk of species range expansion associated with increased temperatures. In Argentina, Aybar et al. [25] found that minimum and maximum temperatures and accumulated rainfall were the best indicators of presence of *C. insignis* and *Culicoides paraensis* Goeldi. Rainfall was also demonstrated as important for the presence of *C. insignis* [26] in Brazil. These studies demonstrate that climate has an effect on the distribution of *Culicoides* spp. and that these data can be used to model the predicted distribution of *Culicoides* species.

Here we modeled the potential distributions of four *Culicoides* species that are putative vectors of orbiviruses in Florida, USA. The four species modeled were *C. insignis, C. stellifer, C. venustus*, and *C. debilipalpis.* The primary objectives to modeling each species were to: 1) compare the major environmental variables predicting the distributions of each species; and 2) produce distribution maps of these species in Florida for use by researchers.

## Methods

### Species Occurrence Records

We built models using two different datasets, including data from the Southeastern Cooperative Wildlife Disease Study (SCWDS), and the Cervidae Health Research Initiative (CHeRI). Data from SCWDS were collected between November 2007 and September 2013 using CDC miniature light traps with white incandescent lights until 2009 and then ultraviolet light emitting diodes (UV/LEDs) from 2009 in order to increase the sampling effort as described in Vigil et al. (2014). CHeRI personnel collected samples from 2015 and June 2017 at deer farms throughout the state as well as state forests and wildlife management areas. 2,910 traps were set by SCWDS over the course of seven years across 59 counties in Florida, and 702 traps were collected by CHeRI in 33 counties from 2016 – 2017 (125 traps used for model validation) and was used to build and validate the models (Figure 1). Only CHeRI data from 2017 were used to validate the models.

**Figure 1.**
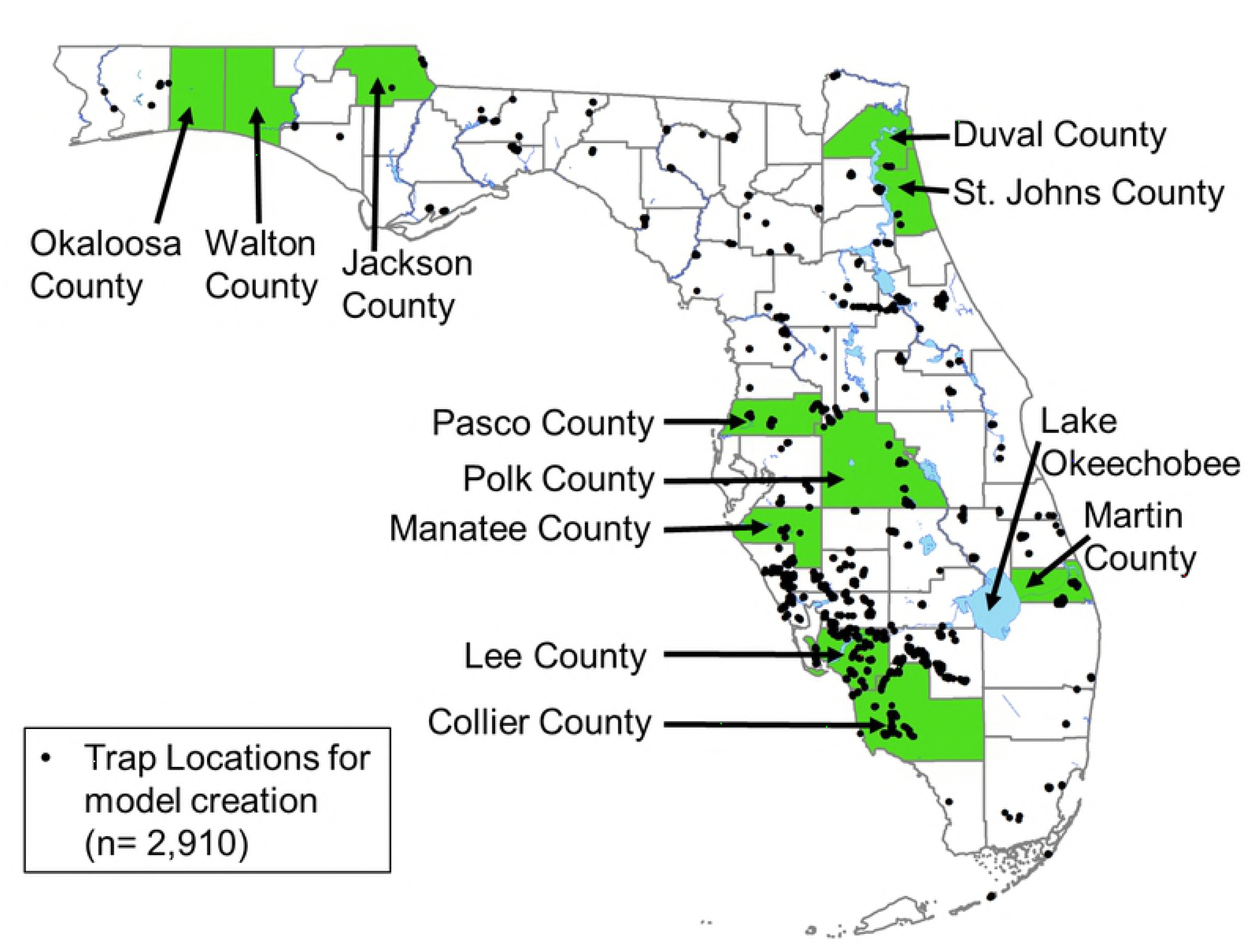
All *Culicoides* spp. trapping locations available for model creation including some county locations. Data were collected by SCWDS and CHeRI between 2008-2017.

We used collection data from the Florida Mosquito Control Association (FMCA), and State Forests during the 2017 field season to validate best model subsets for *C. insignis, C. stellifer*, and *C. debilipalpis.* Participating Mosquito Control Districts (MCDs) included: South Walton County MCD, North Walton Mosquito Control, Jacksonville Mosquito Control (Duval County), Anastasia Mosquito Control (St. Johns County), Pinellas County Mosquito Control, Manatee County MCD, Martin County Mosquito Control, Lee County MCD, and Collier MCD. All CDC miniature light traps were outfitted with UV/LED light arrays and were occasionally supplemented with CO_2_ when available to increase the number of *Culicoides* in the trap. In addition to trapping efforts from county mosquito control districts, permits were obtained to trap on state forest land at the following Florida state forests: John M. Bethea State Forest, Goethe State Forest, Belmore State Forest, Tate’s Hell State Forest, Matanzas State Forest, Blackwater River State Forest, Point Washington State Forest, Lake Talquin State Forest, and Pine Log State Forest. Each state forest was visited between one to three nights throughout the field season, and up to six traps were placed per trapping event.

### Model Covariates

A combination of environmental coverages was used including three bioclimatic variables downloaded from WorldClim (Table 1) (termed “BioClim variables” hereafter), which were derived from the interpolation of historical ground station measures of temperature and precipitation (33). In addition, satellite-derived environmental variables from the Trypanosomiasis and Land Use in Africa (TALA) Research Group (Oxford, United Kingdom) were used describing temperature variable, including middle-infrared (MIR) and land-surface temperature (LST), and vegetation, (NDVI) variables (Table 1) [34]. For this study, we employed a combination of expert opinion and exploratory modeling to determine the final single variable combination that would produce reasonable models for all four species. Our goal was to model each species with the same set of covariates to evaluated potential differences affecting spatial distributions. Pearson correlation tests were used to eliminate highly correlated variables across the nineteen BioClim variables. The final three BioClim variables were chosen based on the limited amount of literature on the ENM modeling of *Culicoides* species [23]. Because vector-borne pathogens cannot exist in the environment without a vector or host, these models can be used to map the potential distribution of disease vectors allowing us to infer disease risk. Other factors such as biotic environmental parameters, genetic diversity, and dispersal may also affect species distributions, however these variables were not modeled as a part of this study [35,36].

**Table 1.** Environmental coverages used for GARP models.

| **Environmental Variable (unit)** | **Name** | **Source** |
| --- | --- | --- |
| **Mean NDVI (no units)** | wd1014a0 | TALA† |
| **NDVI Amplitude (no units)** | wd1014a1 | TALA |
| **NDVI Bi-Annual Amplitude** | wd1014a2 | TALA |
| **Minimum NDVI** | wd1014mn | TALA |
| **Maximum NDVI** | wd1014mx | TALA |
| **NDVI Phase of Annual Cycle** | wd1014p1 | TALA |
| **NDVI Phase of Bi-Annual Cycle** | wd1014p2 | TALA |
| **Mean MIR (°C)** | wd1003a0 | TALA |
| **MIR Annual Amplitude (°C)** | wd1003a1 | TALA |
| **MIR Bi-Annual Amplitude** | wd1003a2 | TALA |
| **Minimum MIR (°C)** | wd1003nm | TALA |
| **Maximum MIR (°C)** | wd1003mx | TALA |
| **MIR Phase of Bi-Annual Cycle** | wd1003p2 | TALA |
| **Mean LST (°C)** | wd1007a0 | TALA |
| **LST Annual Amplitude (°C)** | wd1007a1 | TALA |
| **LST Bi-Annual amplitude (°C)** | wd1007a2 | TALA |
| **Minimum LST (°C)** | wd1007nm | TALA |
| **Maximum LST (°C)** | wd1007mx | TALA |
| **LST Phase of Annual Cycle** | wd1007p1 | TALA |
| **LST Phase of Bi-Annual Cycle** | wd1007p2 | TALA |
| **Max Temperature of Coldest Month** | BIO6 | WorldClim‡ |
| **Precipitation of Wettest Month (mm)** | BIO13 | WorldClim |
| **Precipitation of Driest Month (mm)** | BIO14 | WorldClim |

† Trypanosomiasis and Land Use in Africa (TALA) Research Group (Oxford, United Kingdom)
‡ (www.worldclim.org)
Coverages were obtained from both the Trypanosomiasis and Land Use in Africa (TALA) Research Group, as well as BioClim precipitation and temperature variables from WorldClim.

With every GARP model explained by a genetic algorithm ruleset, we extracted the rulesets of environmental variables [37,38] from the best subsets model output for each species by using GARPTools [29]. For every model, GARP generates 50 rules to predict presence or absence of the species. These rules can be used to evaluate the biological information across models [36,37,39]. The functions in GARPTools automate the extraction of logit and (negated) range rules by (1) identifying and extracting dominant presence rules from the best subsets produced in DesktopGARP, (2) determining the median minimum and maximum values for all environmental variables used for the dominant presence rules in best subset models from the output file in step 1, (3) rescaling all median minimum and maximum values for each variable to 0. 0 – 1.0 to allow for direct comparisons between variables and (4) plotting results in a bar graph to allow for a visual comparison between environmental variables. This process allows for a direct comparison of the differences between variable ranges predicting presence across species.

### Ecological Niche Modeling Approach

Species-specific ENM experiments were run using presence data for the adult locations for each *C. insignis, C. stellifer, C. venustus*, and *C. debilipalpis.* A strategy focusing on precipitation, temperature, and normalized difference vegetation index (NDVI) variables was selected, holding the selected environmental covariates constant for all fours species. All experiments were performed using DesktopGARP v1.1.3 (University of Kansas Center for Research, Lawrence, KS) using the best subset procedure [28]. Presence data were filtered for spatially unique occurrences at 1.0 km × 1.0 km, the native raster resolution of the environmental covariates. Prior to introduction to GARP, unique points were randomly split into testing (25%) and training (75%) datasets for external model accuracy assessment. Experiments used an internal 75% training/25% testing split of occurrence, specifying 200 models at a maximum of 1000 iterations with a convergence limit of 0.01. All four rule types (i.e. range, negated range, logit, and atomic rules) were selected in all experiments. The ten best models in each experiment were chosen using a 10% omission/50% commission threshold [28]. Best subsets were summated into single raster ranging from 0 (absent) to 10 (highest model agreement of presence) [12] illustrating the potential geographic distribution of each species for the entire state. All data preparation for GARP experiments was performed using the GARPTools R-package developed by Haase et al. [29] (IN REVIEW). Once experiments were completed, GARPTools was used to post-process rasters (summation) and calculate accuracy metrics. GARP models are best evaluated using a combination of area under the curve (AUC), omission (false negative rate), and commission (false positive rate; percent of pixels predicted present) [30]. AUC is not an ideal metric [31], but is useful for identifying models that predict well. Final maps were created in ArcGIS v 10.5 (ESRI, Redlands, CA).

*Culicoides venustus* was rarely collected with the trapping method of this study, therefore all available data (CHeRI, SCWDS, MCD, and State Forests) from 2008 to 2017 were pooled from all data sources to build models with adequate sample size. Predictions for *C. venustus* were not validated with field data. Despite the inability to validate the models, building a model for *C. venustus* was still deemed crucial, as evidence points to this species as a potential vector of EHD, based on vector competence studies [32], and its association with host animals during times of outbreak [19]. The preceding work being some of the only published work on this species, it is critical to provide any information we can to the literature.

### Model validation

Field validation points were derived from Mini-CDC light traps (BioQuip Products, Inc.) with black light UV/LED arrays (Model #: 2790V390). When available, carbon dioxide was delivered to the traps via an insulated thermos filled with one gallon of dry ice. The trap netting at the bottom was modified for collecting insects directly into 90% EtOH in a 50mL conical bottom tube. The modification was made by inserting window screen onto the end of the trapping bag normally used for trapping mosquitoes and directing small insects through this screen and into the ethanol-filled tube. *Culicoides* species identifications were based on external morphology of the female using Blanton and Wirth [27].

## Results

Four final experiments were developed for *C. insignis, C. stellifer, C. venustus*, and *C. debilipalpis* (Figure 2). The results of the AUC scores for all models indicated models performed better than random, and most of the models had a total omission of zero, meaning all independent test points were predicted landscapes (Table 2). For the results of the original models made with SCWDS and CHeRI data, the model predicted 96.0% of the presence points of *C. insignis*, across 63.8% of the landscape (Figure 3, Table 2). For *C. stellifer*, the models predicted 96.0% of the presence points across 71.7% of the landscape of Florida, while the models predicted 93.0% of the presence points correctly across 72.5% of the landscape. *Culicoides venustus*, which had the lowest dataset available to build models (n = 18), was most accurate, correctly predicting 98.0% of presence points across just 27.4% of Florida (Figure 3, Table 2).

**Figure 2.**
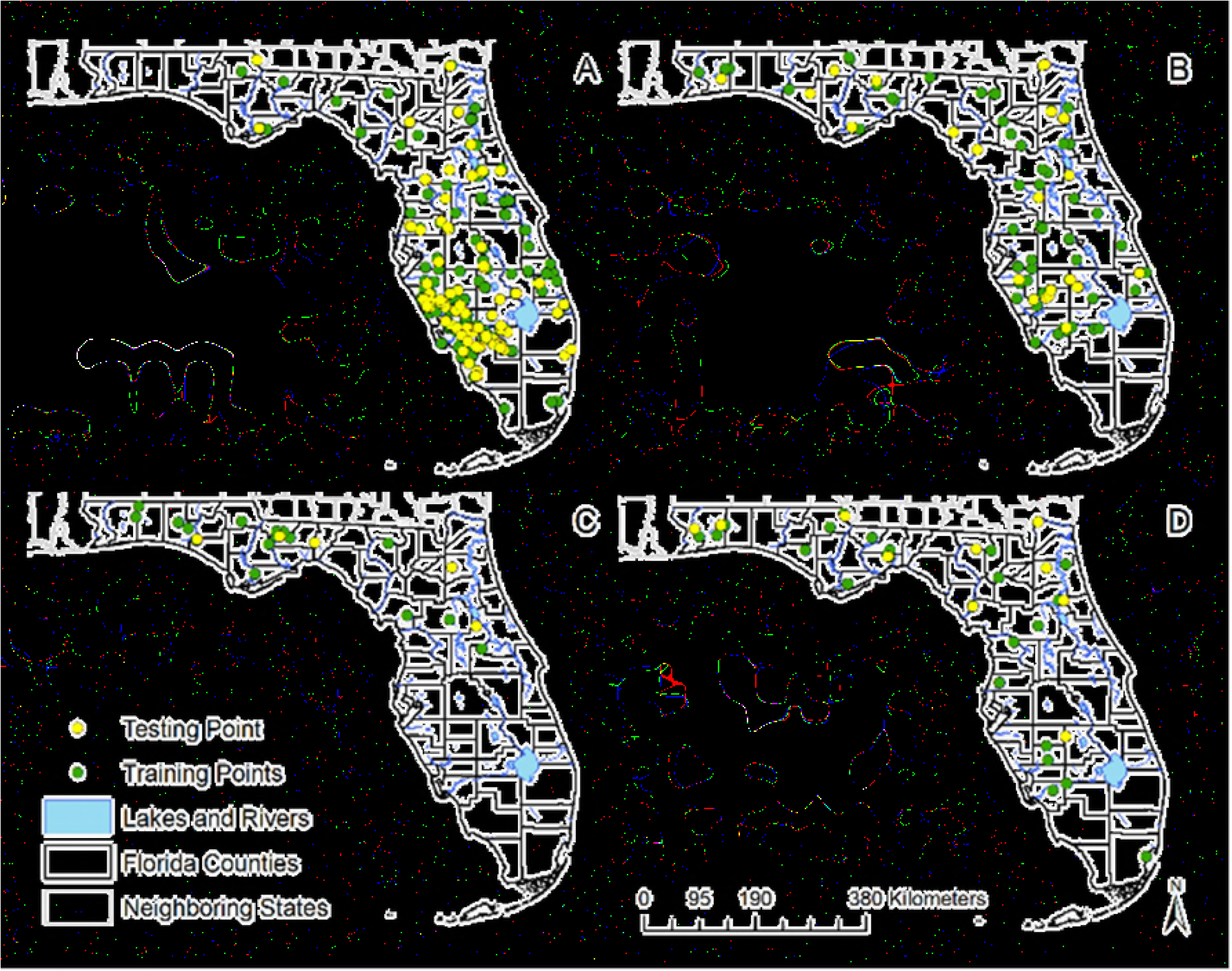
Geographic distribution of the training and testing presence points of four *Culicoides* species of Florida. These species have veterinary importance in Florida and presence points were used in the ecological niche model building and evaluation in desktop GARP. Recorded species distributions in Florida, USA are shown for (A) *C. insignis*, (B) *C. stellifer*, (C) *C. venustus*, and (D) *C. debilipalpis.*

**Figure 3.**
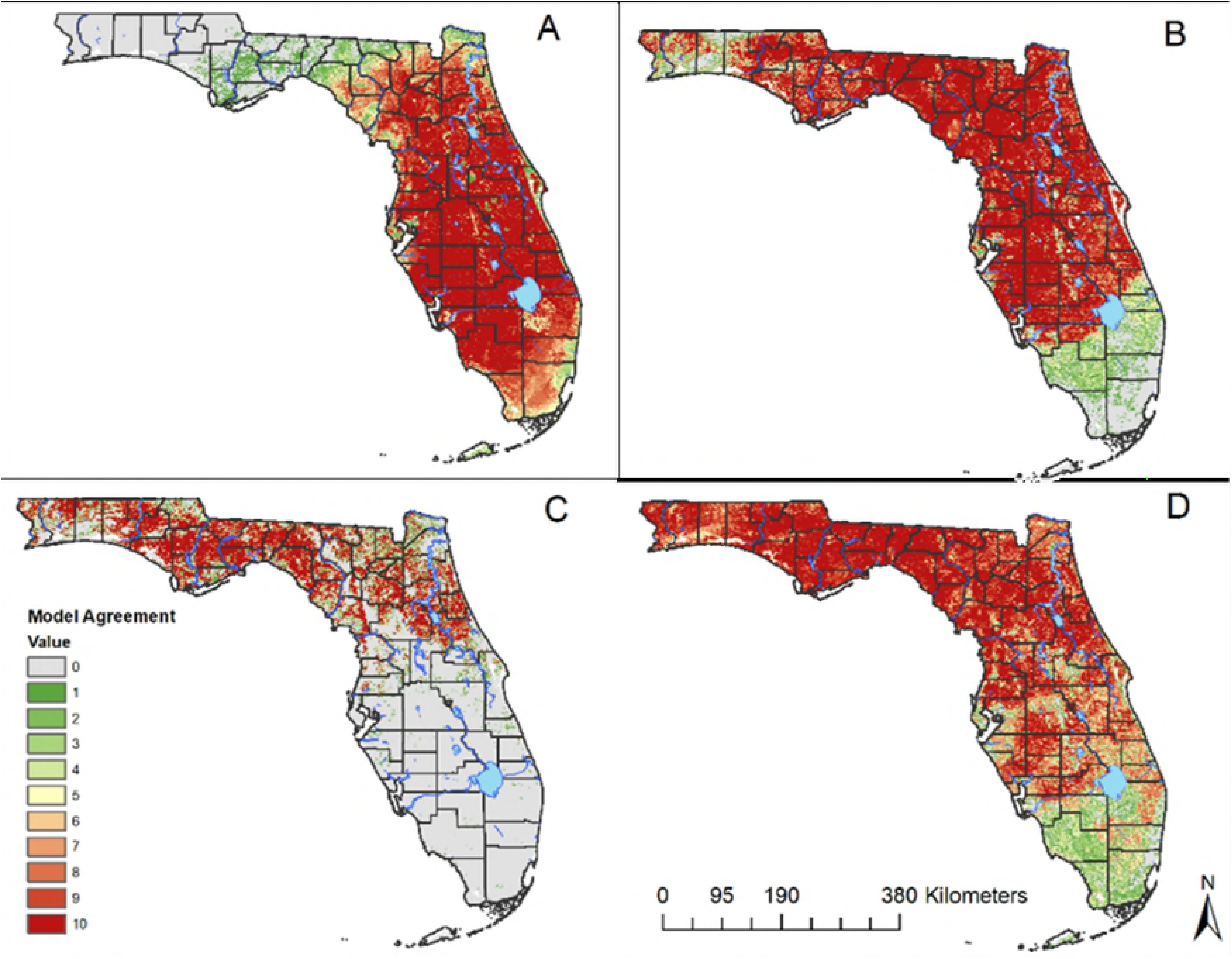
Predicted distributions of *Culicoides insignis* and transferred projections of three other *Culicoides* species. The original model was built for (A) *Culicoides insignis*, (B) *Culicoides stellifer*, (C) *Culicoides venustus*, and (D) *Culicoides debilipalpis*, and models were subsequently transferred for three other species. The color ramp represents the Model Agreement for each species, with zero indicating none of the models predict presence, and 10 all models predict presence.

**Table 2.** Accuracy metrics and sample sizes for GARP model building and evaluation for *C. insignis, C. stellifer, C. venustus*, and *C. debilipalpis.*

| Metric | Model |  |  |  |  |  |  |  |
| --- | --- | --- | --- | --- | --- | --- | --- | --- |
|  | *C. insignis*<br>(CHERI and<br>SCWDS) | *C. insignis*<br>(Validation) | *C. stellifer*<br>(CHERI and<br>SCWDS) | *C. stellifer*<br>(Validation) | *C. venustus*<br>(CHERI and<br>SCWDS) | *C. venustus\**<br>(MCD and<br>State Forest) | *C. debilipalpis*<br>(CHERI and<br>SCWDS) | *C. debilipalpis*<br>(Validation) |
| N to build<br>models | 312 | - | 76 | - | 18 | - | 31 | - |
| N to test<br>models | 104 | 31 | 24 | 23 | 5 | - | 10 | 17 |
| Total<br>Omission | 0.02 | 0.2162 | 0 | 0 | 0 | - | 0 | 0 |
| Average<br>Omission | 3.98 | 34.00 | 4 | 11.74 | 2 | - | 7 | 9.41 |
| Total<br>Commission | 41.17 | - | 53.19 | - | 17.96 | - | 38.53 | - |
| Average Commission | 63.77 | - | 71.66 | - | 27.42 | - | 72.53 | - |
| Standard Error | 0.0280 | 0.0468 | 0.0595 | 0.0624 | 0.0963 | - | 0.0930 | 0.0721 |
| Z-score | 16.9194 | 6.0720 | 8.1300 | 6.4398 | 3.4000 | - | 4.2300 | 5.6337 |
| AUC* | 0.7477 | 0.4768 | 0.6754 | 0.5840 | 0.8891 | - | 0.7000 | 0.6757 |

Also included are the accuracy metrics and sample sizes for the data used for model validations for the GARP models for *C. insignis*, *C. stellifer*, and *C. debilipalpis*. Model for *C. venustus* was not able to be validated.

The *C. insignis* model predicted this species to be widely distributed across the peninsular region of Florida (Figure 3A). The distribution of *C. stellifer* was predicted widely across much of the state, persisting with high model agreement throughout the panhandle south toward Lake Okeechobee in southern Florida (Figure 3B). *Culicoides venustus* had the most geographically restricted prediction of the four species and was predicted to occur primarily in the panhandle and northern Florida. Finally, *C. debilipalpis* was predicted across the largest portions of Florida compared to the other three species modeled. Model agreement was highest in peninsular and northern Florida and model agreement decreased southward (Figure 3D). Disjunct suitable areas were predicted for *C. insignis* into the panhandle region of north Florida (Figure 3A). Predictions for *C. insignis* were also relatively low in the extreme southern part of the state including the Florida Keys, where the landscape is more likely to be dominated by saltmarsh species [40] (Figure 3A). *Culicoides stellifer* was predicted to have low suitability south of Lake Okeechobee (Figure 3B), whereas *C. venustus* was predicted to have no suitable habitat south of Polk County, outside of isolated pixels (Figure 3C).

The experiment for *C. insignis* predicted the 66.0% of 2017 field validation data correctly, while 88.3% of *C. stellifer* locations were predicted correctly and *C. debilipalpis* locations were p with 93.0% accuracy (Table 2). Models were unable to be validated for *C. venustus* due to the relative rarity of this species using our trapping methods and all available data from data sources were used to build models; for *C. venustus*, we relied on the independent testing/training split to assess accuracy (Table 2).

Broadly, covariates with narrow ranges can be interpreted as the most limiting in defining species distributions. Across species, mean, minimum, and maximum LST, and mean, minimum, and maximum MIR had overlapping ranges. Maximum, minimum, bi-annual amplitude and annual amplitude of NDVI were major predictors of *C. venustus* presence, and to a lesser extent, *C. insignis.* Spatial distributions for *C. insignis* were primarily influenced by maximum, minimum and mean LST, as well as maximum, minimum and mean MIR (Figure 4). To a lesser extent, minimum and maximum NDVI predicted presence of *C. insignis*, influencing distributions more than *C. stellifer* and *C. debilipalpis*, but less than *C. venustus.*

**Figure 4.**
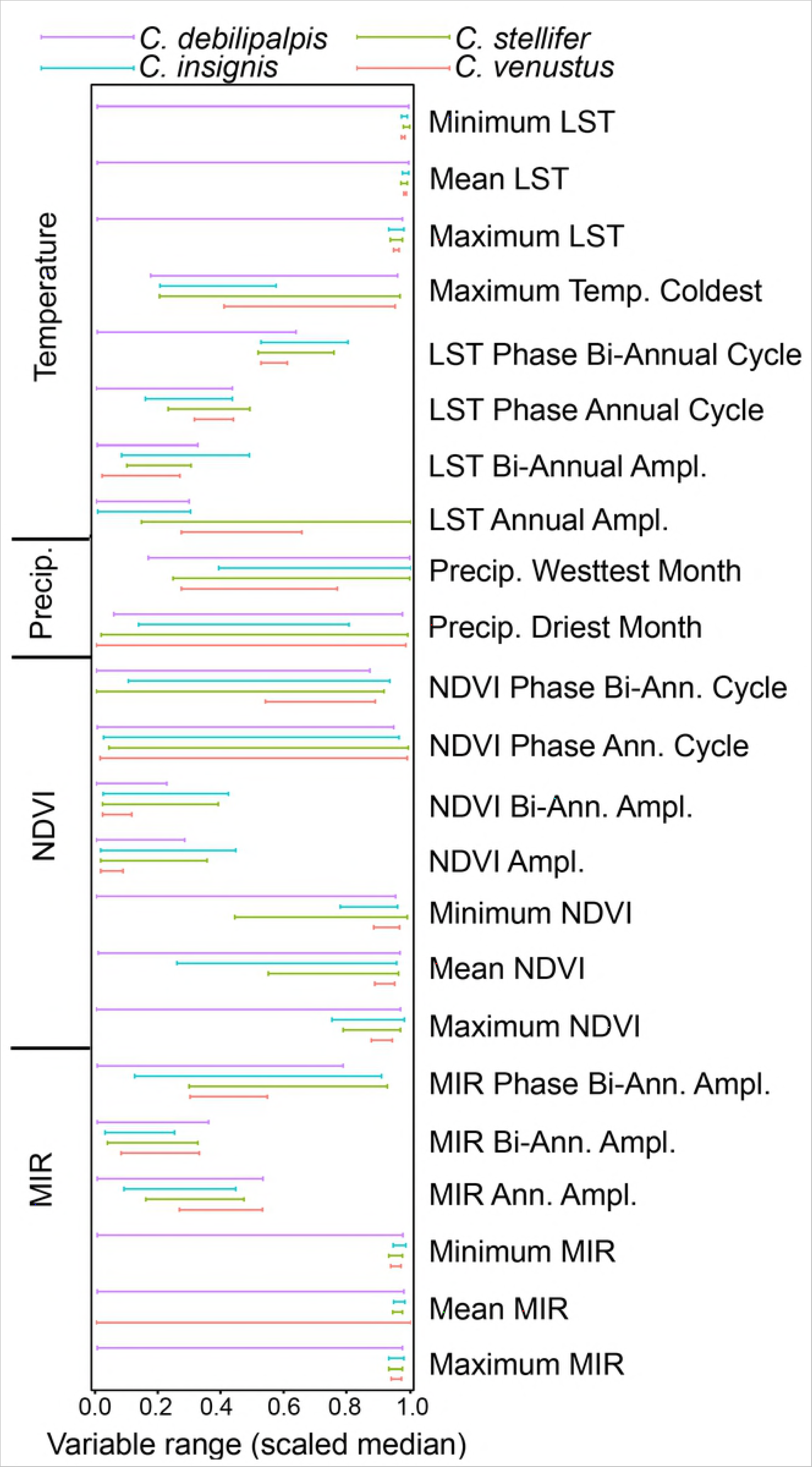
Scales median ranges of the environmental variables for four *Culicoides* species. Ranges are from 0.0 to 1.0 of the environmental variables which predict *C. debilipalpis, C. insignis, C. stellifer and C. venustus* presence.

The median ranges of environmental covariates in the distribution of *C. stellifer* were similar to those of *C. insignis.* For example, minimum, maximum, and mean LST, and maximum, minimum and mean MIR were most limiting for *C. stellifer* (Figure 4). However, unlike *C. insignis*, minimum NDVI was not a limiting covariate for *C. stellifer. Culicoides venustus* had the most limiting covariate ranges of the species modeled in this study. In general, precipitation variables were not found to be important in predicting distributions for any of the four *Culicoides* spp. modeled in this study.

The majority of the environmental covariates used in the model were not sufficient for predicting presence of *C. debilipalpis* as 11 of the 23 selected parameters predicted species presence across more than 90% of their median ranges. The environmental parameters with the lowest percent median range predicting presence were NDVI Bi-Annual Amplitude and NDVI Annual Amplitude at 22.5% and 28.1%, respectively.

## Discussion

This study employed ecological niche modeling to estimate the variable space and geographic distribution of four important *Culicoides* species with veterinary significance in Florida. Though there are several informative studies from Europe, South Africa, and South America using ENMs for *Culicoides* vectors [23,25,41], there is a paucity of this type of work in the Americas.

The distribution for *C. stellifer* was widely predicted across the state with presence unlikely south of Lake Okeechobee. This prediction agrees with historical maps [27] as well as the 2017 field validation data, though efforts to collect in 2017 were limited south of Lake Okeechobee. The low model agreement of *C. stellifer* throughout Okaloosa County (Figure 1) as predicted here (Figure 3B) was also noted in Blanton and Wirth [27] and confirmed with the 2017 field validation data. MIR and LST (both temperature variables) were found to be the most limiting factors for *C. stellifer*, with NDVI being moderately limiting. *Culicoides stellifer* is a widespread, temperate species of *Culicoides*, with its distributions encompassing all of the continental United states and well into Canada [27]. According to the models their presence should be expected to be considerably less likely south of Lake Okeechobee, which marks the Everglades/Lake Okeechobee basin [42] (Figure 3B). This region is the start of a tropical rainy climate, signifying the beginning of a dramatic increase in temperature and precipitation [43], for which *C. stellifer* is probably not as well adapted. Extreme high temperatures and precipitation experienced in the tropical rainy climate of south Florida, in addition to providing a possible upper-limit for heat tolerance of this species, may not provide necessary plant species to contribute to required soil chemistry of *C. stellifer* oviposition sites. Recent studies have shown that *C. stellifer* prefers larval habitats with mud and vegetation from northern Florida over manure or control substrates for oviposition [44]. This should be taken into account when considering the higher probability of the predicted distributions of *C. stellifer* in northern through central Florida, where temperate plant species are abundant [45].

The models for *C. debilipalpis* also predicted the species across much of Florida. However, this prediction is somewhat counter-intuitive to the data collected during the present study; it was not very common using our sampling method. This result may be attributed to the inability of our light traps to attract all species of *Culicoides* equally [46]. It is possible that *C. debilipalpis* is present in the areas not predicted by the model.

The large areas absent of *C. debilipalpis* could also be explained by habitat preference. *Culicoides debilipalpis* has been confirmed to develop in tree-holes of *Salix* spp. [47]. Although the specific larval habitat species of *C. debilipalpis* have not yet been elucidated in Florida, they are probably also located in tree-holes of several different hardwood species. For this reason, the *C. debilipalpis* map outputs, driven by NDVI, may show areas where fewer hardwood species are available for larval *C. debilipalpis* development in the green and grey portions of the map (Figure 3D). South of lake Okeechobee, the landscape is dominated by extensive marsh with scattered pine rockland forests and tropical hardwood hammocks [45], habitats that are not known for tree holes.

The predicted distribution of *C. venustus* was consistent with Blanton and Wirth [27]. In North America, distributions for *C. venustus* extend north to Canada and as far west as Oklahoma [27,48]. In this study, *C. venustus* was not recorded further south than Pasco County [27], and the models predict low probability of *C. venustus* south of Pasco County (Figure 3). NDVI variables, in particular, bi-annual cycle, bi-annual amplitude, amplitude, minimum, maximum and mean of the NDVI, were most limiting for *C. venustus* out of all four species of *Culicoides* modeled in this study. In addition, temperatures, particularly the LST, were limiting for the distributions of *C. venustus. Culicoides venustus* is predominantly a northern species [27], making its cold-tolerance higher than other more tropical species, such as *C. insignis.* Similarly, EHDV is most common in deer and *Culicoides* spp. in northern Florida and the panhandle in the same region of Florida where *C. venustus* is most common. This significant overlap between a putative vector species and disease warrants further investigation of this species as a vector of EHDV.

In contrast to both the historical data from Blanton and Wirth [27] restricting *C. insignis* East of Gilchrist Counties, as well as the original models, model validation suggest that *C. insignis*, is likely to occur outside of predicted ranges and into peninsular Florida. *Culicoides insignis* was discovered in the continental United States and Florida in 1948 [49], and was assumed to be restricted to the state until at least 1979 [27]. Hagan and Wirth [50] documented *C. insignis* in Glynn and Chatham Counties, Georgia. We collected *C. insignis* extensively in the panhandle in 2017 and it has been collected in light traps in both Mississippi and Alabama as recently as 2014 [50,51]. Recent studies by Vigil et al. [52] have also documented the range of *C. insignis* extending into Mississippi and Louisiana.

The GARP experiments using SCWDS and CHeRI data predicted *C. insignis* distributions to extend as far west as Jackson County in Florida. CHeRI sampling covered a similar span of longitudes as the SCWDS sampling efforts from 2007-2013, suggesting that *C. insignis* may have been absent from much of the panhandle during the time period the SCWDS data were collected. The low AUC (<0.50) for the validation model from the 2017 validation data suggests that models for *C. insignis* predicted the panhandle poorly.

One possible explanation for poor model prediction in the panhandle may be attributed to an increase in sampling efforts in 2017. State forests in the panhandle were visited several times during the season and we also consistently sampled in North and South Walton Counties. Poor model agreement could also be an indication that *C. insignis* is recently expanding to habitats north of its historic distribution, as Vigil et al., [52] has noted. In other studies, vector *Culicoides* spp. have been documented to have northward movements across continents, including the expansion of *C. imicola* into Europe [41,53,54]. A similar pattern was also observed in *C. sonorensis* by Zuliani et al. [24]. In that study, they used a MAXENT approach to predict the future distributions *of C. sonorensis* in North America. The models predicted that the northern latitudinal limits of the distribution of *C. sonorensis* was at high risk for expansion due to climate change. Similarly, climate change may contribute to the northwest expansion of *C. insignis*, in part, due to findings of the model that *C. insignis* were primarily influenced by high temperature (LST and MIR) variables, which included minimum and maximum annual temperatures which were also found to be significant to their distributions in South America (25). Potential northward expansion of *C. insignis* will likely affect the distributions of endemic BTV infection in cattle and white-tailed deer as it will begin to overlap more with the distributions of another major vector of BTV in North America, *C. sonorensis* [18]. Further, spread of a subtropical vector of BTV could increase the chance of introducing exotic BTV serotypes from South America, as serotypes 1, 3, 6, 8, 12, and 14 have historically been restricted to South America in the New World [55]. This scenario is demonstrated to be plausible by the first report of bluetongue virus serotype 1 in Louisiana in 2006 [56].

A narrow median range suggests that the ecological niche of the species is constrained to the specific conditions of this covariate, which indicates the environmental variable plays a more important role in the distribution of a species [37,38]. Two different measures of average temperature, MIR and LST, were most limiting in predicting *C. insignis.* Additionally, a variable with narrow median range for *C. insignis* was the maximum temperature of the coldest month (Figure 4). *Culicoides insignis* is a tropical species primarily distributed throughout South America where it is the primary vector for BTV [55]. Its range extends to southern Florida, due to the subtropical nature of the climate, where it has been implicated in BTV transmission [21]. As *C. insignis* is primarily a peninsular species, the demonstration of these variables to apparently drive the niche for *C. insignis* suggests that its proclivity to peninsular Florida is driven by temperature. In previous studies, *C. insignis* distribution was influenced by mean minimum humidity [57], which also associates the species with peninsular Florida. Although humidity was not a parameter used in the present study, temperature associations with other *Culicoides* species have been demonstrated previously. *Culicoides imicola*, a non-native *Culicoides* species in Europe, was also demonstrated to have a narrow temperature preference similar to *C. insignis* [58].

Kline and Greiner [59] observed that larval *C. insignis* is found in close association with cattle. This not only provides evidence that *C. insignis* is associated with hosts, it also demonstrates that certain mud characteristics may be more suitable for larval development. This association would indicate that soil pH in particular could be used to make the models stronger *for C. insignis*.

Overall, NDVI Bi-Annual Amplitude, NDVI Amplitude, MIR Bi-Annual Amplitude, MIR Annual Amplitude, as well as LST Phase of Annual Cycle and LST Bi-Annual Amplitude were limiting variables across all four *Culicoides* species analyzed in this study (Figure 4). None of the chosen environmental parameters appear to meaningfully predict *C. debilipalpis* along a narrow median range. For *C. debilipalpis*, the environmental covariates with the narrowest median ranges predicting presence were NDVI Bi-Annual Amplitude and NDVI Annual Amplitude. The other covariates were less useful to predict *C. debilipalpis* and future work should focus on modeling each species with the most appropriate covariates rather than a uniform set for all four species.

Environmental variable ranges were much narrower for *C. insignis, C. stellifer*, and *C. venustus*, suggesting a higher confidence that variables used in this study are driving distributions of some species. NDVI was a limiting factor for all four species of *Culicoides.* As these *Culicoides* spp. breed in semi-aquatic soils, the chemical composition of which is influenced by surrounding plant species [60], which can potentially change the measurement of the NDVI [61]; the NDVI median range is potentially specific to each species of *Culicoides* [47,62].

In conclusion, all four *Culicoides* species demonstrated unique distributions using the chosen environmental variables in GARP experiments. *Culicoides insignis* was predicted widely in the peninsular region, *C. stellifer* was predicted north of Lake Okeechobee, *C. venustus* was predicted in the panhandle and north, and C. *debilipalpis* was predicted across the largest range of Florida, although with large gaps in predicted presence in between. Minimum, and maximum MIR and minimum, maximum, and mean LST were the most limiting variables for *C. insignis, C. stellifer*, and *C. venustus*, indicating that upper and lower temperature limits are most important in the distributions of these *Culicoides* species. In contrast, these were some of the least limiting environmental variables used for *C. venustus*, for which the bi-annual and annual amplitudes of LST, MIR, and NDVI were more limiting. Most NDVI variables were also very limiting for the distributions of *C. venustus* and were moderately limiting for *C. debilipalpis.* In general, however, the chosen environmental variable set was not suitable for predicting the presence of *C. debilipalpis.* These ranges suggest that temperature (LST and MIR) is the most limiting factor in the distributions of these *Culicoides* spp., followed by NDVI. In general, precipitation variables were not found to be important in predicting distributions of these four *Culicoides* spp. in Florida.

This study employed ecological niche modeling to estimate the variable space and geographic distribution of four important *Culicoides* species with veterinary significance in Florida, marking the first of its type in Florida and is among very few distribution modeling studies for *Culicoides* species in North America. It should be noted that our focus for the model building was primarily on exploring the differences in environmental conditions and holding these variables constant between four different species in the genus *Culicoides*. This approach allowed for the direct comparison between the median ranges of environmental variable. As a result, these maps may not be the most geographically accurate maps for each species. Future work should focus on modeling each species with the most appropriate covariates rather than a uniform set, modeling the ideal covariates for each species in order to create more accurate distribution maps. Accurate maps will further our understanding of EHDV and BTV transmission and may be used by wildlife managers (spanning public and private lands), and livestock producers to identify areas most at risk for EHDV and BTV disease transmission. Further, identification of environmental covariates contributing to their distributions will allow for the development of an integrated pest management program to control vector species.

## Acknowledgements

The authors wish to thank the following persons at mosquito control districts (MCD) for their help with trapping in 2017: P. Brabant and B. Brewer (South Walton County MCD), V. Hoover, G. Hubbard, B. Hunt, G. Morse, and D. Wallace (North Walton Mosquito Control), M. Clark (Jacksonville Mosquito Control), D. Dixon, and R-D Xue (Anastasia Mosquito Control), B. Bayer, and E. Buckner (Manatee County MCD), K. Zirbel (Martin County Mosquito Control), J. Stuck (Pinellas County MCD), R. Morreale (Lee County MCD), and M. Kartzinel and R. Bales (Collier MCD). Florida Department of Agriculture and Consumer Services, and State Forest Ecologist Brian Camposano for administering the permit to trap in state forests. The authors also wish to thank H. Lynn, and A. Runkel for help with sorting and identification, E. Blosser for their assistance with fieldwork, A. Quaglia for assistance with Figure 4, and G. Ross for their technical assistance. J. P. Gomez and C. Lippi provided assistance with niche modeling data preparation. This work was supported by the University of Florida Institute of Food and Agricultural Sciences, Cervidae Health Research Initiative.

## Supporting Information

**S1 Table.** Coordinates of each presence point used to build the final models for each of the four species used in this study: *C. insignis, C. stellifer, C. debilipalpis*, and *C. venustus.* These coordinates are adjusted from the original dataset to remove duplicates in the same raster. As such the dataset is spatially unique at 1.0km.

